# Fentanyl reinstatement to discriminative cues after conflict in sign- and goal-tracking rats

**DOI:** 10.1101/2022.02.10.479913

**Authors:** David A. Martin, Sara E. Keefer, Donna J. Calu

## Abstract

**Rationale:** Discriminative stimuli (DS) are cues that predict reward availability. DS are resistant to extinction and motivate drug seeking even after long periods of abstinence. Previous studies have demonstrated that sign tracking (ST) and goal tracking (GT) individual differences in Pavlovian approach to food cues predict distinct vulnerabilities to CS and DS reinstatement of cocaine seeking, respectively. Compared to goal-trackers, sign-trackers show heightened CS relapse even after electric barrier induced abstinence. We do not know whether DS relapse persists after electric barrier induced abstinence, or whether tracking-related relapse vulnerabilities generalize to models of opioid relapse.

**Objectives:** We sought to determine if DS-induced reinstatement of fentanyl seeking persists in the presence of reduced adverse consequences after electric barrier-induced abstinence. We also aimed to determine whether tracking differences predict the magnitude of DS-induced reinstatement of fentanyl seeking after electric barrier-induced abstinence.

**Methods:** First we used Pavlovian lever autoshaping (PLA) training to determine sign-, goal- and intermediate tracking groups in male and female Sprague Dawley rats. We then trained rats in a DS model of intermittent fentanyl self-administration, and extinguished drug seeking by imposing an electric barrier of increasing intensity. We then measured the level of DS-induced reinstatement in the presence of a reduced electric barrier intensity.

**Results:** We report that DS produce large increases in fentanyl seeking after electric barrier induced abstinence. Contrary to our expectations, the magnitude of the DS induced reinstatement effect was not related to tracking group.

**Conclusions:** Discriminative stimuli powerfully motivate opioid seeking, despite continued aversive consequences. Individual differences in Pavlovian approach do not predict the level of DS reinstatement to fentanyl seeking after conflict induced abstinence.

## Introduction

Conditioned stimuli (CS) motivate reward seeking behavior when animals learn their association with unconditioned stimuli (US) such as food or drugs. Individual reactivity to food-associated CS predicts drug seeking behaviors in models of addiction vulnerability (Flagel et al. 2010; Saunders and Robinson 2010). Sign tracking (ST) and goal tracking (GT) individual differences in Pavlovian CS approach behavior predict distinct relapse vulnerabilities for cocaine seeking (Saunders and Robinson 2010; Saunders et al. 2014), but this is not the case for remifentanil seeking (Chang et al. 2022). While reinstatement to contingent CS induced drug seeking persists despite negative consequences (Saunders et al. 2013), it is unclear if discriminative stimuli (DS), which predict US availability, also promote reinstatement despite negative consequences. Here, we determine whether DS-induced relapse to fentanyl seeking persists following conflict induced abstinence and whether ST and GT behaviors predict the magnitude of reinstatement effects.

The temporal relationships of CS relative to US delivery determine how stimuli affect behavior (Di Ciano and Everitt 2003). Discriminative stimuli (DS) predict when a reward seeking response will produce the US, whereas contingent CS are present only after a reward is earned. Both DS and contingent CS stimulate reward seeking, but they function through different mechanisms (Di Ciano and Everitt 2003; Namba et al. 2018). DS function similarly to contexts, informing when rewards are available, whereas contingent CS function as conditioned reinforcers. DS paired with cocaine resist extinction and promote drug seeking that escalates with abstinence (Weiss et al. 2001; Madangopal et al. 2019). In humans, DS may be important drivers of relapse to drug seeking, as exposure to cues predictive of drug availability occurs before relapse – whereas contingent interoceptive and environmental stimuli associated with drug taking affect behavior after relapse has occurred. Therefore, defining the conditions that influence DS control of drug-seeking behavior will increase our understanding of drug relapse. The present study aims to determine whether DS predictive of opioid availability trigger relapse after negative consequences are imposed.

Sign- and goal-tracking individual differences in approach toward a food-associated CS correlate with differences in behaviors motivated by drug-associated CS and DS in relapse models (Saunders and Robinson 2010; Saunders et al. 2013, 2014). In Pavlovian lever autoshaping (PLA) procedures, a lever cue predicting food motivates either lever pressing, termed sign-tracking (ST), or food-magazine exploration, termed goal-tracking (GT). Previous work shows that individuals that preferentially exhibit ST to a food cue are more prone to both food (Yager and Robinson 2010) and cocaine (Saunders and Robinson 2010) reinstatement when a discrete CS is paired with delivery of reward during training, whereas GT individuals are more susceptible to contextual and DS-induced reinstatement of cocaine seeking (Saunders et al. 2014; Pitchers et al. 2017).

These relationships support the hypothesis that individual differences in incentive salience attribution to CS predict reinstatement of reward seeking across diverse US (cocaine and food), and that the type of cue (predictive DS vs. contingent CS) may dictate whether ST or GT rats are more susceptible to reinstatement. However, it remains unknown whether patterns of reinstatement vulnerability to predictive cues (DS) are stable across other US types, such as for opioids.

The conflict aspect of the model we employ in this study uses a footshock barrier of escalating intensity to extinguish drug taking (Cooper et al. 2007), followed by reinstatement of DS-induced drug seeking in the presence of reduced footshock barrier intensity. The conflict induced abstinence phase is designed to model the aversive consequences of drug seeking and drug taking that fluctuate in humans. Here, we examine whether DS-induced reinstatement to fentanyl seeking persists after conflict induced abstinence and whether ST and GT predict sensitivity to DS-induced relapse to opioid seeking despite conflict. Based on prior studies with psychostimulants (Pitchers et al. 2017), we predicted that GT individuals would exhibit greater DS-induced reinstatement of fentanyl seeking even in the presence of the shock barrier.

## Materials and Methods

### Subjects

Male and female Sprague-Dawley rats (Charles Rivers Laboratories, Wilmington, MA; 200-275 g upon arrival; n=56 run as 2 separate cohorts) were 8 weeks old and triple housed with same-sex cagemates upon arrival. Rats were maintained on a 12 hr reverse light/dark cycle (lights off at 10:00 am), and all behavioral training and testing were conducted during the dark phase of the cycle. Rats had *ad libitum* access to standard laboratory chow and water throughout all phases of the experiment. Rats were single housed after acclimation and prior to behavioral training. All behavioral experiments were performed in accordance to the “ Guide for the Care and Use of Laboratory Animals” (8^th^ edition, 2011, US National Research Council) and were approved by the University of Maryland, School of Medicine Institutional Animal Care and Use Committee (IACUC).

### Catheterization Surgery

After establishing tracking phenotype with Pavlovian lever autoshaping (see below), we anesthetized rats with isoflurane (5% induction, 1-3% maintenance) and implanted catheters into the right jugular vein. The catheter was made from Silastic tubing (cat#508-002, Dow Silicones Corp, Midland, MI, USA), subcutaneously inserted, and affixed to the 22-gauge guide stainless steel backmount cannula (PlasticsOne, Roanoke, VA, USA) that protruded through a small back incision. We subcutaneously administered a non-steroid, anti-inflammatory drug, carprofen (5 mg/kg, Rimadyl ®) prior to surgery and for three days after surgery. We infused rats daily with 0.05 mL of anti-microbial, anti-bacterial, and anti-coagulant Taurolidine-Citrate (TCS) catheter lock solution i.v. (Cat# TCS-04, Access Technologies, IL, United States) to reduce biofilm and clot formation, to promote catheter patency, and to reduce the risk of microbial infection throughout the experiment. We checked catheter patency periodically via i.v. injections of 0.1 mL of methohexital sodium (“ Brevital”), and rats without a sudden loss of muscle tone were removed from the study (n=8).

### Drugs

We purchased fentanyl citrate from Cayman Chemical and diluted it in 0.9% sterile saline to 1 mg/mL before further diluting in 20 ml syringes to concentrations scaled to each rat’s weight for a dose of 1ug/kg/injection.

### Behavioral Procedures

Experimental design is outlined in Figure 1A. Behavioral experiments were conducted in identical behavioral chambers (25 ⨯ 27 ⨯ 30 cm; Med Associates) located in rooms different than the colony room. Each chamber was located in individual sound-attenuating cubicle with a ventilation fan. Each chamber had a red house light (6 W) located at the top of the wall opposite the experimental stimuli.

**Fig. 1:**
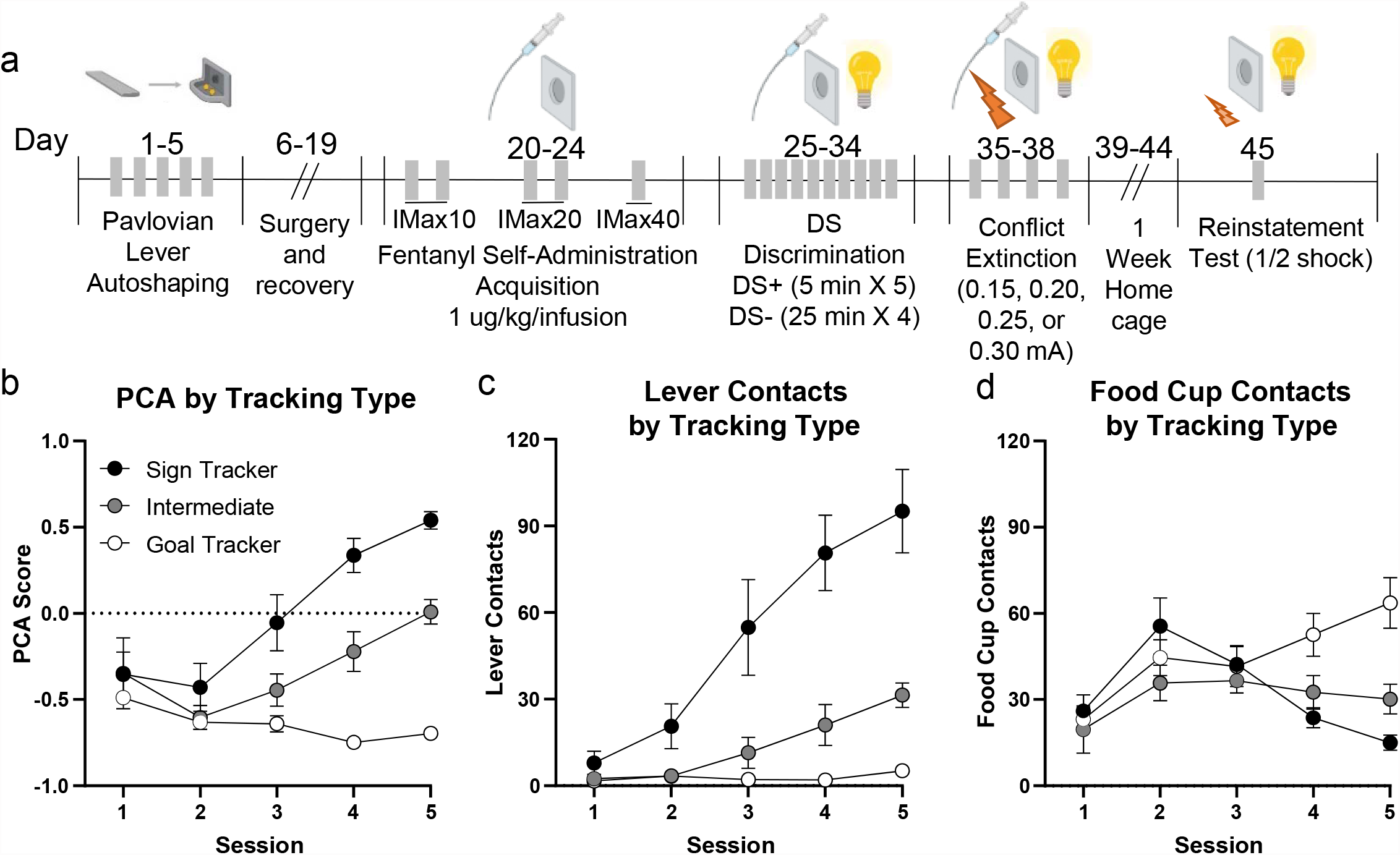
A) Experimental timeline B-D) Pavlovian conditioned approach training: B) Overall PCA tracking score C) Lever contacts D) Food cup contacts

## Pavlovian lever autoshaping

### Apparatus

For Pavlovian lever autoshaping, the red house light was illuminated for the duration of each session. The opposite wall had a recessed food cup located 2 cm above the grid floor. The food cup had an attached programmed pellet dispenser to deliver 45 mg food pellets (catalog#1811155; Test Diet Purified Rodent Tablet (5TUL); protein 20.6%, fat 12.7%, carbohydrate 66.7%). A retractable lever was positioned on either side of the food cup 6 cm above the floor, and side was counterbalanced between subjects.

### Behavioral Procedure

We habituated rats to the food pellets prior to training. Then, we trained rats for five daily, ∼26 min Pavlovian lever autoshaping (PLA) sessions. Each session included 25 presentations of a lever presentation that served as the conditioned stimulus (CS) and occurred on a VI 60 s schedule (50-70s). The lever was inserted for 10 s for each trial, retracted, and followed immediately with the delivery of two food pellets into the food cup. Food delivery occurred independent of lever or food cup approach or contact. After each training session, we transported rats back to the colony room.

### Behavioral Measurements

We used a Pavlovian Conditioned Approach (PCA) analysis (Meyer et al. 2012) to determine sign- and goal-tracking groups. PCA quantifies the continuum of lever-directed (sign-tracking, ST) and food cup-directed (goal-tracking, GT) behaviors. A PCA score is calculated for each rat and is the average of three difference score measures, including 1) preference score, 2) latency score, and 3) probability score, each ranging from - 1.0 to +1.0. For preference score, we recorded total number of contacts with the lever and food cup during the CS. The calculation of preference score was the number of lever contacts during the CS minus the number of food cup contacts during the CS, divided by the sum of these two measures. Latency to first contact to lever and food cup during the CS was recorded, and if contact did not occur, a latency of 10 s was recorded. The calculation of latency score was the average latency to make a food cup contact during the CS minus the latency to lever contact during the CS, divided by the duration of the CS (10 s). Lever and food cup probabilities were calculated by determining the number of trials that the response was made divided by total number of trials in the session. The calculation for the probability score was the probability of a lever contact minus the probability of a food cup contact throughout the session. We averaged the PCA scores during session 5 to determine tracking groups. Sign-tracking PCA scores range from +0.33 to +1.0, goal-tracking PCA scores range from -0.33 to -1.0, and intermediate PCA scores range from -0.32 to +0.32.

## Fentanyl self-administration, discriminative training, conflict, and relapse

### Apparatus

In a separate room from PLA training, we trained rats in self-administration chambers contained in sound attenuating cabinets (Med Associates) similar to PLA training, but stimuli were different from PLA to not confound actions between trainings. One wall contained two nose pokes located 5 cm above the grid floor with a white light located between them 10 cm above the grid floor. A red light was located at the top of the wall on the opposite side. We counterbalanced the side of the Active nosepokes for each tracking group relative to the side of the lever during PLA training.

### Self-administration Acquisition and Discriminative Stimulus Training

After a 7-12 day recovery from catheterization surgery, we trained rats in 5 sessions to self-administer fentanyl for 2 h per session. A nose poke into the active poke activated the syringe pump to deliver 1 ug/kg fentanyl in 28 ul over 1 second on a fixed ratio 1 schedule. A nose poke into the inactive poke was recorded, but no fentanyl was delivered. No stimuli were paired with nosepokes or drug infusions, and no lights were turned on during initial training. As in prior tracking studies (Saunders et al. 2013; Pitchers et al. 2017), we imposed an infusion maximum (IMax) capping the maximum number of infusions/session to limit differences in acquisition of self-administration between tracking groups.The IMax was 10 infusions/session for two sessions (IMax10), 20 for two sessions (IMax20), and 40 for one session (IMax40). We concluded each rats’ session either when reaching IMax or at 2 hours from session start, whichever came first. The IMax10 and IMax20 sessions used a 20 s timeout following each infusion during which active nosepokes were recorded but did not result in additional infusions. The IMax40 session and all subsequent discriminative stimulus (DS) sessions used a 1 second timeout corresponding to the length of the infusion.

After 5 IMax self-administration acquisition sessions, we trained rats in 10 sessions using an intermittent access (IntA) schedule with two distinct discriminative stimuli signaling drug availability (DS+) or non-availability (DS-). These sessions began with 2 min illumination of the red house light (DS-) followed by a 5 min illumination of a white light (DS+) located on the opposite wall between the two nose pokes. During the DS+, a response into the active nose poke resulted in delivery of fentanyl on a fixed ratio 1 schedule with 1 second timeout. We imposed a limit of seven infusions per 5 min DS+ to avoid the potential for overdose. After the 5 min DS+, the white light turned off, and the DS-red light illuminated for 25 min signaling drug was unavailable. This pattern was repeated 3 more times before the session ended following a final DS+ period. The total session length was 127 minutes, consisting of five, 5-minute DS+ periods, and four, 25-minute DS-periods between them, in addition to the first, 2-minute DS-period. We recorded all active and inactive nose pokes during the session.

### Conflict

We imposed a conflict-induced abstinence model that introduced a negative consequence of increasing footshock intensity to decrease drug-seeking and taking behaviors while the reinforced DS schedule maintained. Sessions were similar to IntA sessions with DS+ and DS-, except an electric current was constantly applied to the two-thirds of the grid floor closest to the nose pokes throughout the entire session. As a result, rats had to traverse the electric grid to nose poke and receive fentanyl infusions. We trained rats in four daily sessions of conflict IntA sessions, with the footshock intensity set to either 0.15, 0.20, 0.25, or 0.30 mA using an aversive stimulator (Med Associates). All rats started at 0.15mA, and if they received more than 5 infusions at this shock intensity, we increased shock intensity by 0.05 mA the following session. If they received less than 5 infusions, we repeated the same shock intensity the following session. Rats remained in their colony room for a week after the fourth day of conflict prior to reinstatement test.

### Reinstatement Test

After one week in their home cages with no testing, we tested rats in a reinstatement test under extinction (no drug available) conditions, although the animals were still tethered to a drug delivery line. Similar to conflict training, the two-thirds of the grid floor closest to the nose pokes were electrified, but to 50% of each rat’s maximum shock intensity reached during conflict training, consistent with a prior tracking study investigating CS-induced relapse after conflict-induced abstinence (Saunders et al. 2013). DS+ and DS-periods were shorter in duration than during training (30 sec and 150 sec, respectively), but the ratio of DS+/DS- durations was identical. After an initial 2 min DS- to begin the session, the DS+ was on for 30 s followed by the DS-for 150 s. The session ended after 21 DS-/DS+ cycles (62.5 min total). Nose pokes into the active and inactive ports were recorded.

### Statistical Analysis

One, two, and three way ANOVAs were employed with post-hoc tests corrected for multiple comparisons where appropriate. Repeated measures were employed appropriately for mixed within-subject/between subject designs. Sphericity corrections were performed where appropriate with Greenhouse-Geiser corrections. Pearson’s correlation analysis was performed to correlate PCA scores with reinstatement behavior. Tests were performed in Prism (Graph Pad). All data shown is that from rats that completed the entire study (n=28). The remainder were either not surgerized following PLA testing (n=12), lost catheter patency before finishing the study (n=8), failed to reliably self-administer fentanyl (>5 infusions/session; n=3), or became sick and were removed the study (n=5). We calculated a discrimination score as a measure of discrimination between DS+ and DS- response rates using the following equation:

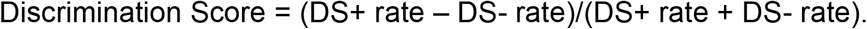

## Results

### Acquisition of Pavlovian Autoshaping

Prior to any drug experience, we screened male and female rats using PLA to classify their tracking phenotype as sign-tracking (ST), goal-tracking (GT), or intermediate (INT). Pavlovian conditioned approach (PCA) scores serve as a comprehensive index summarizing the number of contacts, latency to contact and probability to contact the lever or food cup across the session (Fig. 1A). We classified rats based on their performance on session 5 of training (see Methods for details). We analyzed the PCA scores over the five days using a mixed-design, repeated measures ANOVA, with a between-subject factor of Tracking Group (ST (n=7), GT (n=16), INT (n=5)) and a within-subject factor of Session (Fig. 1B). Based on Session 5 characterization we observed an interaction between Tracking Group and Session (F_(8,100)_ = 16.54, p<0.0001), indicating that behavior motivated by a food-predictive lever stimulus developed differently between the assigned tracking groups, as expected (Fig. 1B). The differences in PCA scores are characterized by increases in pressing across sessions in STs, but not GTs (Fig. 1C), and an increase in pokes in GTs, but not STs (Fig. 1D).

### Acquisition of Fentanyl Self Administration

After determination of tracking groups, we implanted intravenous jugular catheters in ST, GT and INT rats. After recovery we trained rats to nosepoke for fentanyl infusions (1 ug/kg/infusion) in five 2 hour sessions. To ensure there were no tracking related differences in the initial acquisition of fentanyl self-administration, we imposed an infusion maximum (IMax) capping the total number of infusions/session to 10, 20, or 40. No cues were explicitly paired with drug infusions, although the activation of the syringe pumps is audible. All rats included in the study reliably discriminated active from inactive nosepokes (Fig. 2A), as shown by a main effect of Nosepoke (active vs inactive) (F_(1,54)_=19, p<0.0001) and an interaction between Nosepoke and Session (F_(4, 108)_ = 4.334, p=0.0027). Active poking increased between session 1 and session 5 (p=0.0007, Dunnett’s test), whereas inactive poking remained similar between session 1 and 5 (p=0.9438, Dunnett’s test).

**Fig. 2:**
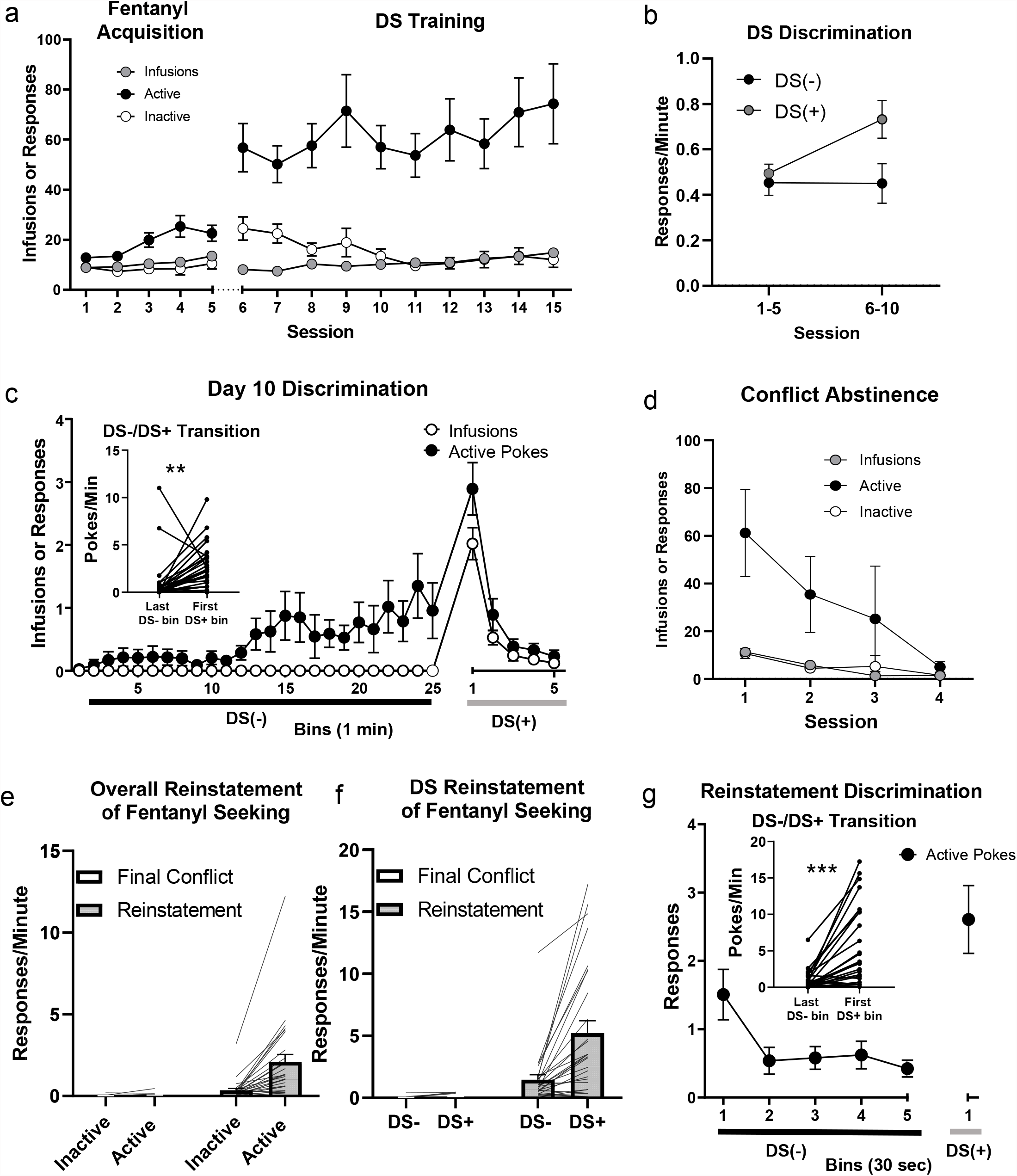
Fentanyl motivated behavior for all rats combined. A) Fentanyl self-administration through acquisition (five sessions) and DS training phases (ten sessions). B) Rates of responding in the DS+ and DS- components of DS training sessions, binned by the first five and last five sessions of DS discrimination phase. C) Day 10 of DS training. Data are binned into 1 min segments across the 25 min DS- component and 5 min DS+ component, and the resulting data are averaged across all DS-/DS+ cycles across the session. Inset shows the individual data for the transition between the last bin of the DS- component and the first bin of the DS+ component (**: p<0.01, paired t-test). D) Conflict extinction behavior in the presence of electrified floor barrier in front of nosepokes. E) Poking rates (inactive/active) for the last conflict extinction test (left two columns) and reinstatement test (right two columns). F) Poking rates (DS-/DS+) for the last conflict extinction test (left two columns) and reinstatement test (right two columns). G) Reinstatement test poking rate data binned into 30 s segments across the 2.5 min DS- component and 30 s DS+ component. Data are averaged across all DS- /DS+ cycles in session. Inset shows the individual data for the transition between the last bin of the DS- component and the first (only) bin of the DS+ component (***: p<0.001, paired t-test).

### Discriminative Stimuli Training

After initial fentanyl self-administration training, we introduced discriminative stimuli to distinguish drug available (ON) from non-available (OFF) periods. The DS+ (signaling ON periods) was a white light in between the two nosepokes, and the DS- (signaling OFF periods) was a red light on the back wall. Overall, rats maintained active and inactive nosepoke discrimination throughout DS training (main effect of Nosepoke: F_(1,27)_=45.32,p<0.0001), and increased the number of infusions received across DS sessions (one-way ANOVA; effect of Session: F_(4.67, 126.2)_=10.56, p<0.0001) (Fig. 2A). Importantly, rats reliably learned to discriminate DS+ and DS- stimuli over the course of the ten DS sessions (Fig. 2B). This learning is partially captured by higher DS+ active poking rates relative to DS- rates overall (main effect of Stimulus (DS+,DS-): F_(1, 27)_=13.56, p=0.001), as well as by a Stimulus (DS+,DS-) x Session interaction (F_(1, 27)=_15.92, p<0.001).

To further investigate how rats responded to the discriminative stimuli, we examined behavior on a finer timescale by separately binning DS- (25 min) and DS+ (5 min) periods into 1 min bins and averaging across all DS-/DS+ cycles within a DS session. In DS session 1, before rats have learned the stimuli, we observe a relatively uniform distribution of responding over time across the DS-/DS+ cycle (Fig. S1A). In particular, we observe no difference between the last minute of responding during the DS- period and the first minute of responding during the DS+ period (Fig. S2A, inset). These two minutes capture the transition from DS- OFF to DS+ ON periods, when the white light turns on to signal fentanyl availability. In contrast, in DS Session 10, responding slowly increases across the DS- period before suddenly increasing in the first minute of DS+ periods (p=0.003, paired t-test; Fig. 2C, inset). These data indicate that by the final DS session, fentanyl seeking behavior was under the control of the DS schedule.

### Conflict Induced Abstinence of Fentanyl Responding

Following ten sessions of DS training, we maintained the DS schedule of fentanyl reinforcement but imposed an electrified barrier in front of the nosepoke apparatus. We applied constant footshock to the front 2/3 of the floor grid, such that rats had to move across the electrified floor to reach the nosepokes. Consistent with a prior conflict relapse study investigating tracking differences (Saunders et al. 2013), we increased footshock intensity incrementally across 4 sessions (0.15, 0.20, 0.25, or 0.30 mA) if the number of infusions earned in the prior session did not drop below 5. If the number of infusions earned dropped below 5, shock intensity remained constant for the next session(s). We observed a marked decrease in infusions earned across the four sessions (RM one-way ANOVA, effect of Session: F_(2.64, 71.3)_=24.43, p<0.0001) as well as a decrease in all responses (main effect of Session: F_(3,81)_=5.076, p=0.0029) (Fig 2D).

### DS+ Induced Reinstatement of Responding

Rats spent one week in their home cages with no testing prior to a DS reinstatement test. For this test we used 1/2 maximum shock intensity reached for each rat during conflict-induced abstinence, consistent with prior CS conflict relapse study (Saunders et al. 2013). We tested rats under extinction conditions (no drug available). Overall, active responding rates dramatically increased during reinstatement testing compared with the final day of conflict induced abstinence (Fig. 2E). We observed significant main effects of Response (active vs. inactive) (F_(1, 27)_ = 23.27, p <0.0001) and Session (Conflict induced abstinence vs. Relapse test) (F_(1, 27)_ = 18.28, p =0.002), as well as a significant interaction between Response and Session (F _(1,27)_=21.50, p<0.0001). During the reinstatement test, active poking rate during the DS+ periods was significantly elevated relative to the final day of conflict-induced abstinence (Fig. 2F). We observed main effects of Stimulus (DS+,DS-) (F_(1, 27)_ = 21.29, p <0.0001) and Session (F _(1, 27)_ = 26.06, p <0.001), as well as a Stimulus x Session interaction (F (_1,27)_=19.99, p=0.0001). We observed a >50 fold increase in active nosepoking rate and >65 fold increase DS+ poking rate in the Reinstatement vs. Final Conflict Session (Fig 2E-2F). These data indicate that a reduction in conflict magnitude led to a large increase in drug seeking responses controlled by the DS schedule of reinforcement.

To further investigate the pattern of DS responding, we analyzed the Reinstatement Test data by binning DS- periods (150 sec) and DS+ periods (30 sec) into 30 sec bins, and averaging binned poking rates across all 21 equivalent DS-/DS+ cycles (Fig. 2G). Similar to our Day 10 training data, we observed an increase in active pokes between the last DS- bin and the first DS+ bin (p<0.0001, paired t-test) (Fig 2G, inset). These data indicate that discriminative stimuli associated with fentanyl availability powerfully motivate drug seeking following conflict-induced abstinence in the face of attenuated but continued conflict.

#### Tracking and Sex as Factors

To assess tracking type and sex as factors during DS discrimination training, conflict induced abstinence, and DS-induced reinstatement of fentanyl seeking, we compared responding across all experimental phases, independently considering tracking type and sex as factors. The statistics for these comparisons are summarized in Table 1.

**Table 1:** All statistical comparisons for measures compared between tracking types and sexes.

| Main Effects | Acquisition Infusions | Acquisition Active Pokes | DS Infusions | DS Active Pokes | DS Discrimination | Conflict Infusions | Conflict Active Pokes | Conflict Shock Intensity | Last Conflict vs Reinstatement Active Pokes | Last Conflict vs Reinstatement DS+ Pokes |
| --- | --- | --- | --- | --- | --- | --- | --- | --- | --- | --- |
| Tracking | F(2,25) = 0.602<br>P=0.555 | F(2, 25) = 2.076<br>P=0.146 | F(2, 25) = 1.21<br>P=0.3131 | F(2, 25) = 0.609<br>P=0.5518 | F(2, 25) = 0.202<br>P=0.8183 | F(2, 25) = 0.488<br>P=0.6191 | F(2, 25) = 1.108<br>P=0.3459 | F(2, 25) = 0.843<br>P=0.4420 | F(2, 25) = 1.949<br>P=0.1635 | F(2, 25) = 0.756<br>P=0.4796 |
| Sex | F(1,26) = 0.409<br>P=0.5280 | F(1, 26) = 0.075<br>P=0.7863 | F(1, 26) = 0.502<br>P=0.4846 | F(1, 26) = 0.300<br>P=0.5881 | F(1, 26) = 0.803<br>P=0.3782 | F(1, 26) = 3.433<br>P=0.0753 | F(1, 26) = 0.914<br>P=0.3478 | F(1, 26) = 1.349<br>P=0.2560 | F(1, 26) = 0.305<br>P=0.5851 | F(1, 26) = 0.300<br>P=0.5884 |
| Interactions |  |  |  |  |  |  |  |  |  |  |
| Tracking by Session | F(8,100) = 1.57<br>P=0.1434 | F(8, 100) = 1.33<br>P=0.2354 | (18, 225) = 0.737<br>P=0.7699 | F(18, 225) = 0.848<br>P=0.6413 | F(2, 25) = 0.059<br>P=0.9425 | F(6, 75) = 0.947<br>P=0.4667 | F(6, 75) = 1.145<br>P=0.3452 | F(6, 75) = 0.704<br>P=0.6473 | F(2, 25) = 1.899<br>P=0.1707 | F(2, 25) = 0.704<br>P=0.5040 |
| Sex by Session | F(4,104) = 1.189<br>P=0.3200 | F(4, 104) = 1.179<br>P=0.3246 | F(9, 234) = 1.378<br>P=0.1991 | F(9, 234) = 0.862<br>P=0.5599 | F(1, 26) = 1.014<br>P=0.3232 | F(3, 78) = 0.331<br>P=0.8025 | F(3, 78) = 0.569<br>P=0.6366 | F(3, 78) = 1.266<br>P=0.2920 | F(1, 26) = 0.378<br>P=0.5436 | F(1, 26) = 0.234<br>P=0.6326 |

During Acquisition and DS training we found no differences in infusions earned between tracking groups and no Tracking and Session interactions (Fig 3A). Similarly, we found no main effects of Tracking or interactions between Tracking and Session on active poking rates during acquisition or during DS training (Fig S2B).

**Fig. 3:**
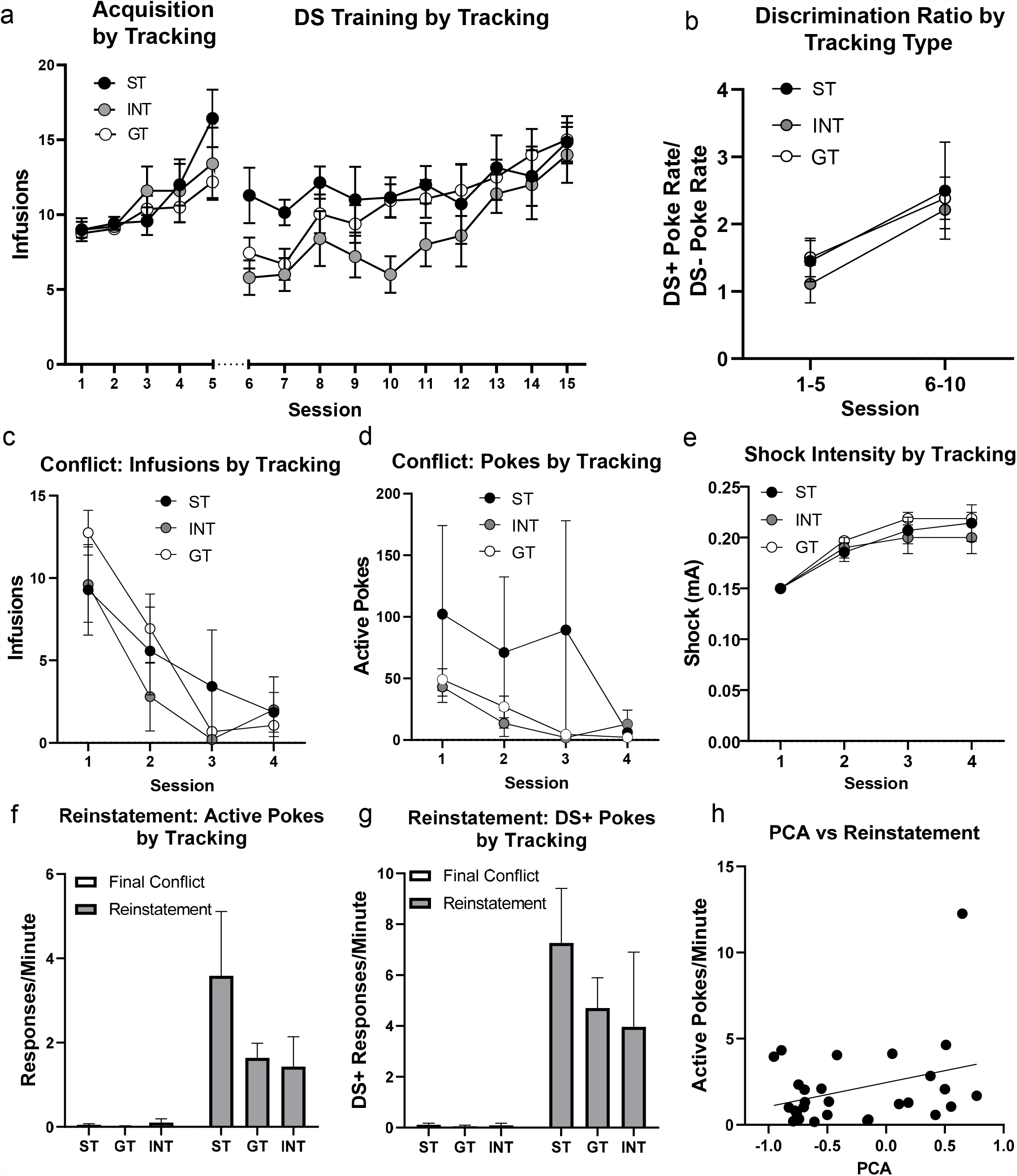
Fentanyl motivated behavior between groups defined by tracking type. A) Infusions of fentanyl through acquisition and DS training phases. B) Discrimination ratio during the first five sessions and last five sessions of DS training. C) Conflict extinction infusions D) Conflict extinction active pokes E) Average shock intensity used across sessions F) Active side poking rates during final conflict session (left) and reinstatement test (right). G) DS+ poking rates during final conflict session (left) and reinstatement test (right). H) Linear correlation between PCA score and active poking rate. (R2=.1023, p=0.0971).

Collapsing across tracking groups to compare the sexes, we found no differences in acquisition response rates and infusions, nor in DS training response rates or infusions, with respect to sex (Fig S1A, S2C). These data indicate that neither tracking nor sex were associated with differences in acquisition of fentanyl self-administration, which may in part be due to the IMax schedule we imposed. Furthermore, by the end of DS training, all tracking groups and both sexes were nearly identical in average number of infusions earned per session (Fig 3A, S1A).

To understand potential differences between tracking groups in DS discrimination, we compared the ratio of DS+/DS- responding between early (Sessions 1-5) and late (Sessions 6-10) DS Training (Fig. 3B). While we found a main effect of Session, we found no main effects of Tracking, nor a Tracking by Session interaction, indicating that all three groups’ relative ratios of DS+/DS- responding increased similarly across training (Fig 3B). Analogously, to understand potential differences between sexes in DS discrimination, we compared the ratio of DS+/DS- responding between early and late sessions and found no significant effects of Sex nor any interaction between Sex and Session (Fig S1B). Taking into account that the DS-:DS+ transition is best captured by the rate of responding between the last DS- bin and first DS+ bin of DS-/DS+ cycles (see above), we compared these transitions between tracking types using a discrimination score (see Methods), and again found no significant differences in discrimination score between groups on the last day of training (Fig S2D), indicating all tracking groups expressed DS discrimination similarly.

As conflict training progressed, all tracking groups reduced their infusions earned (Fig 3C) and active responding (Fig 3D) across sessions and we found no main effects of Tracking nor interactions of Tracking with Session during conflict-induced abstinence. We also compared the increases in shock intensities across tracking groups (Fig. 3E), and again found no main effects of Tracking, nor any significant interactions. Analogously, we compared across sexes for infusions (Fig S1C), active responding (Fig S1D), and shock intensities (Fig S1E). Again, we found no main effects of Sex, nor any interactions of Sex with Session for any of these three measures. These data indicate all tracking groups and both sexes were similarly sensitive to conflict-induced abstinence of fentanyl seeking.

In order to examine whether tracking group influenced DS+ reinstatement after conflict induced abstinence, we compared the active nosepoke response rates (Fig. 3F) and DS+ response rates (Fig. 3G) between tracking groups and across the final conflict session and the reinstatement test session. Although we found a large main effect of Session, we found no main effects of Tracking, nor a Session by Tracking interaction (see Table 1 for statistics). Analogously, we compared active response rates (Fig S1F) and DS+ response rates (Fig. S1G) between the sexes and across the final conflict session and reinstatement session. Again, we found no main effects of Sex, nor any Sex by Session interactions on these measures. We also compared DS discrimination between tracking groups by again calculating a discrimination score between the last DS- bin and first DS+ bin during reinstatement, and found no difference between tracking groups on this measure (Fig S2D). Finally, we examined whether there was any linear relationship between PCA score and active poking rate during reinstatement and found no significant relationship (R2 = .102, p = .0971)(Fig 3H). The weak trend for a larger magnitude reinstatement effect in sign-tracking rats was mostly driven by a single rat (see Fig 3H). These data suggest that although the DS+ serves as a powerful fentanyl reinstatement stimulus in the face of continued but reduced conflict this effect does not vary by tracking type or sex.

## Discussion

We first classified rats as ST, GT, or INT by their Pavlovian approach behaviors and then trained them to nosepoke to self-administer fentanyl. We then introduced an intermittent access schedule in which ON periods were signaled by a DS+, and OFF periods were scheduled by a DS-. Over the ten days of DS training, rats learned the relationship between the DS and drug availability, as evidenced by a sudden ramping of drug seeking behavior at the onset of the light DS+. Under conditions of conflict between escalating shock intensity and continued fentanyl reinforcement, all rats reduced their drug seeking and intake to very low levels as shock intensity increased. In accordance with our predictions, following a week of homecage abstinence, we observed robust DS+ controlled reinstatement under extinction conditions with reduced-intensity (1/2 maximum shock intensity). Additionally, we compared the development of DS+ controlled drug taking, conflict-induced abstinence, and DS+ induced reinstatement across ST, GT, and INT rats. Throughout all stages of our experiments, and contrary to our expectations, we found no significant differences between rats across tracking groups. We observed no sex differences in these same behaviors between male and female rats.

Our experimental conditions combine conflict-induced abstinence designs (Cooper et al. 2007; Saunders et al. 2013) that model adverse consequences associated with human drug use and DS designs, which model environmental cues signaling whether drugs are available. Our primary goal in employing this design was to determine if DS+ stimuli are sufficient to produce robust reinstatement of opioid seeking following conflict induced abstinence under conditions of continued, but reduced, conflict. Indeed, we observed very high rates of drug-seeking behavior during the reinstatement test, and this drug seeking was highly concentrated in the DS+ periods. Many studies have shown that DS+ stimuli powerfully modulate reward seeking responses (eg. Dinsmoor 1950; McFarland and Ettenberg 1997). Drug-associated DS in particular motivate behavior that is resistant to extinction (Martin-Fardon and Weiss 2017) and increases with the passage of time (Madangopal et al. 2019). Our results here add to previous results by showing that DS+ associated with opioid availability motivate vigorous drug seeking behavior in the face of continued conflict.

In addition to establishing robust DS+ reinstatement of fentanyl seeking under conflict, we sought to compare and correlate the intensity of reinstatement across individuals with differences in PCA behavior. Numerous studies have found that sign- and goal-tracking behaviors in response to a food cue predict the intensity of cue-induced reinstatement of drug taking – in a cue-dependent fashion (Robinson et al. 2014). While sign-tracking correlates with discrete contingent cue-induced reinstatement (Saunders and Robinson 2010), goal-tracking has been associated with contextual reinstatement (Saunders et al. 2014) and DS+ reinstatement (Pitchers et al. 2017). Directly relevant to our study, recent work has demonstrated a significantly greater DS+ mediated reinstatement of cocaine-seeking in goal-tracking relative to sign-tracking rats (Pitchers et al. 2017). Therefore, the lack of difference between tracking groups in our work is surprising and may be due to one or a combination several differences in the studies, which we explore below.

Importantly, our study used the opioid agonist, fentanyl, as the US whereas the previous study (Pitchers et al. 2017) used the psychostimulant, cocaine. Many studies to date linking PCA behaviors to differential drug-related behaviors have used cocaine as a reinforcer. In addition to established relationships of PCA behavior to cocaine reinstatement (Saunders and Robinson 2010; Saunders et al. 2014; Pitchers et al. 2017), sign-tracking predicts choice of cocaine over food (Tunstall and Kearns 2015), sensitivity to cocaine psychomotor sensitization (Flagel et al. 2008), and acquisition of cocaine self-administration (Beckmann et al. 2011). Additionally, sign-tracking predicts contingent CS-induced reinstatement to methamphetamine (Everett et al. 2020) and nicotine (Versaggi et al. 2016). However, a recent study using the opioid remifentanil found no difference in contingent CS-induced reinstatement of drug seeking between ST and GT rats (Chang et al. 2022). These results are consistent with the present findings for DS-relapse to opioid seeking, which when considered together suggest that drug class (psychostimulant versus opioid) may be an important factor in determining cue-induced reinstatement across tracking groups.

Neither our results nor the results of Chang et al. (2022) were predicted; as heroin (Peters and De Vries 2014), remifentanil (Yager et al. 2015), and cocaine (Yager and Robinson 2013) similarly support approach to Pavlovian drug-paired cues. Pavlovian drug-associated cue approach is magnified in individuals that ST to a food-predictive lever (Yager and Robinson 2013; Yager et al. 2015). However, the inability of PCA behavior to predict DS and CS mediated reinstatement suggests that Pavlovian sign-tracking and reinstatement of opioid seeking in an operant context are unrelated processes (Chang et al. 2022). Interestingly, when using a lever as the Pavlovian cue predictive of drug reward, cocaine supports lever approach (Uslaner et al. 2006), but only heroin supports significant lever pressing (ie. sign-tracking) (Madsen and Ahmed 2015). We further note that PCA behavior to food cues may not correlate in every case to PCA behavior using different US, as previous work showed that sign-tracking for liquid sucrose showed no relationship to sign-tracking for food (Patitucci et al. 2016), and that sign-tracking for food was not related to visual nicotine cue approach (Yager and Robinson 2015). Clearly, further investigation into the importance of the US in the development of PCA behavior is warranted to understand the relationship between PCA behavior and reinstatement using diverse US.

In addition to the US used, our study also differs methodologically with earlier work. To extinguish drug-seeking responses, we used conflict conditions in the presence of continued drug availability under the DS schedule. In contrast, the previous DS+ study removed the drug and DS stimuli during classic extinction procedures in which drug was not available (Pitchers et al. 2017). While in both cases, drug seeking is similarly decreased, the associative processes underlying decreased responding are fundamentally different. Therefore, it is conceivable that differences in reinstatement between tracking groups emerged due to the number or nature of extinction sessions. However, there were no differences in behavior reduction during extinction between tracking groups in either study (Pitchers et al. 2017). Alternatively, different sensitivities between tracking groups to shock might explain the discrepancy, but we observed no group differences in sensitivity to shock during conflict training, consistent with earlier work using this procedure (Saunders et al. 2013). However, others have found greater resistance to punishment in sign tracking animals (Pohořalá et al. 2021).

Our study also employed different conditions for DS training (fewer and shorter sessions) and reinstatement testing than similar previous DS work with cocaine (Pitchers et al. 2017). Perhaps the most notable difference is our DS+ periods during reinstatement were 30 s, followed by an ITI of 150 s; their DS+ presentations were shorter: 4 s followed by an average ITI of 30 s. We observed a very large increase in drug seeking rates across all tracking groups during DS+ vs. DS- periods relative to the more modest DS+ rate increases observed in the prior study (Pitchers et al. 2017). This may potentially be due to the length of our DS+, but the continued presence of conflict during the reinstatement test is also likely to have discouraged responding during DS- periods in our study. Overall, because the discrimination between DS+ and DS- was both robust and equivalent between tracking groups in training and reinstatement in our study, it is difficult to reconcile the marked discrepancy in tracking effects between studies based on differences in DS training and testing parameters.

In the current work, intermittent access to fentanyl was available for many (19) sessions prior to the reinstatement test. We would expect this schedule to considerably increase economic demand for fentanyl over baseline levels (Martin et al. 2020), conceivably strengthening the DS-US relationship across all tracking types as the drug becomes more valuable. Supporting this idea, sign tracking individuals exhibit higher cocaine demand early in training, but extended cocaine experience equalizes this difference, as well as equalizing early differences in cue-induced reinstatement (Kawa et al. 2016). Furthermore, a recent study using 45 sessions found that a composite addiction measure combining persistence of cocaine seeking, motivation for cocaine taking, and resistance to punishment, did not correlate with PCA behavior (Pohořalá et al. 2021). Further work exploring the importance of the drug and amount drug experience in modifying cue-motivated behaviors across different drugs and stimulus types is necessary to understand the complex relationships between traits and experience that drive drug seeking behaviors.

In summary, we report that discriminative stimuli signaling the availability of fentanyl powerfully motivate drug seeking even in the presence of conflict, and the strength of this drug seeking does not correlate with Pavlovian tracking phenotypes. This work adds to growing literature delineating the complex relationship between sign- and goal-tracking behaviors and addiction models.

## Acknowlegements

This work was funded through National Institute on Drug Abuse (NIDA) RO1 grant # RO1DA043533 awarded to Donna Calu.

---

Figure Captions for: Fentanyl reinstatement to discriminative cues after conflict in sign- and goal-tracking rats.

**Fig. S1:**
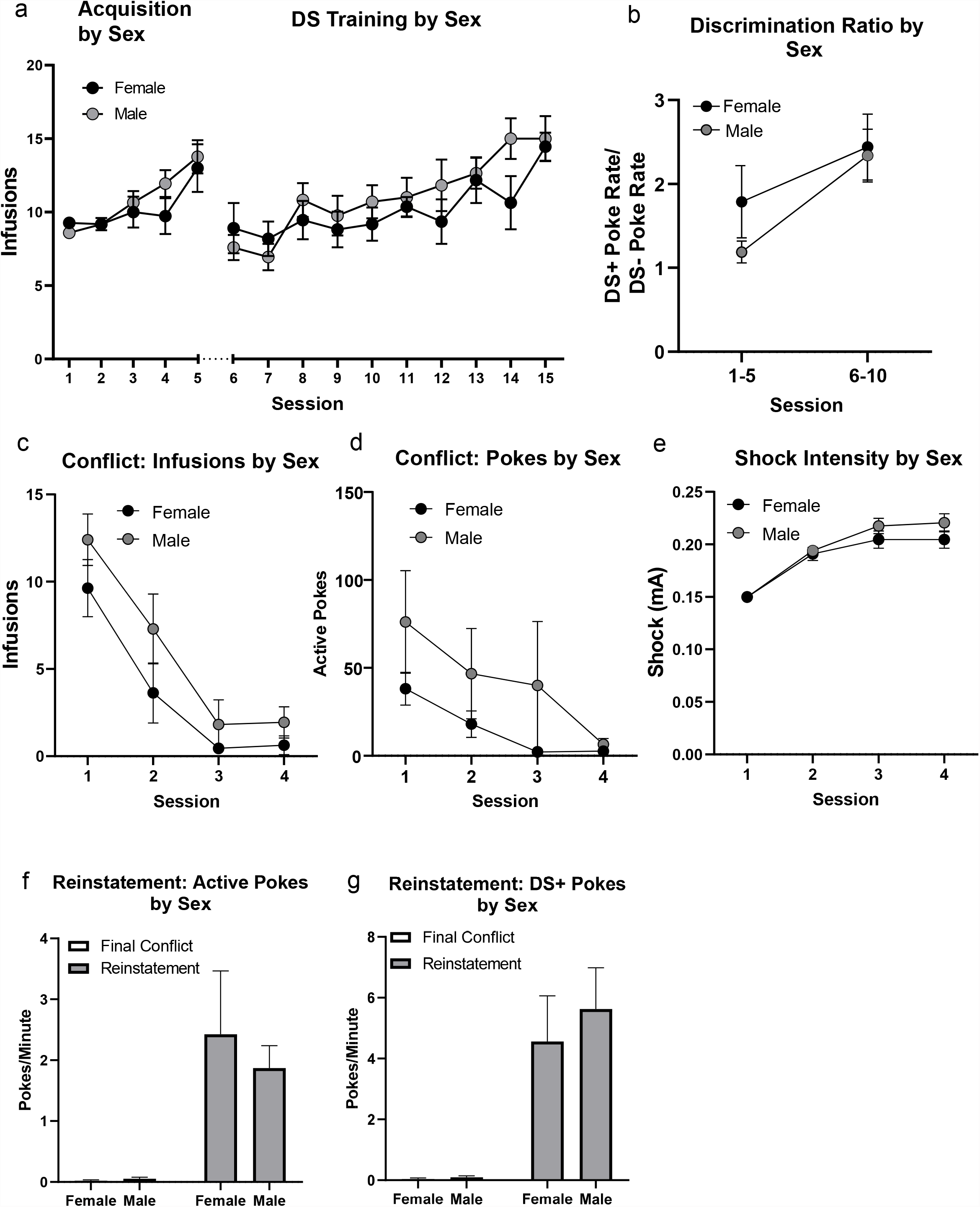
Fentanyl motivated behavior between groups defined by sex. A) Infusions of fentanyl through acquisition and DS training phases. B) Discrimination ratio during the first five sessions and last five sessions of DS training. C) Conflict extinction infusions. D) Conflict extinction active pokes. E) Average shock intensity used across sessions. F) Active side poking rates during final conflict session (left) and reinstatement test (right). G) DS+ poking rates during final conflict session (left) and reinstatement test (right).

**Fig. S2:**
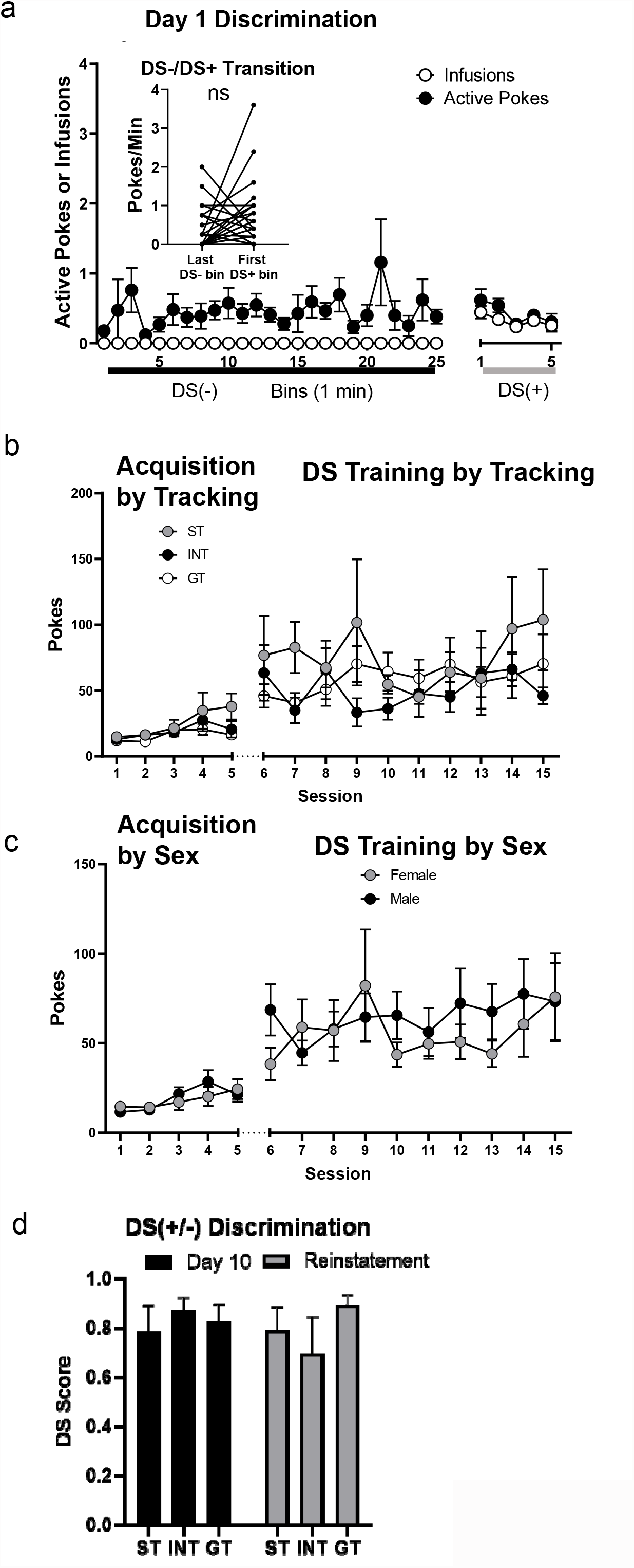
A) Day 1 of DS training. Data are binned into 1 min segments across the 25 min DS- component and 5 min DS+ component, and the resulting data are averaged across all DS-/DS+ cycles across the session. Inset shows the individual data for the transition between the last bin of the DS- component and the first bin of the DS+ component. B) Comparison of tracking groups for active pokes during fentanyl acquisition and DS training phases. C) Comparison of sexes for active pokes during fentanyl acquisition and DS training phases D) Comparison of discrimination scores between tracking groups on Day 10 of DS training and during reinstatement test.

